# Predicting protease networks through human genetics

**DOI:** 10.1101/2022.06.30.498364

**Authors:** Kazunari Iwamoto, Tore Eriksson

## Abstract

By utilizing functional genetic variation within the participants of the UK Biobank project for a largescale PheWAS study we attempted to get a better understanding of how the set of human proteases and their endogenous inhibitors are involved in common diseases. Focusing on known human proteases, their inhibitors, and known substrates, we computed their ranked-biased similarity from phenome-wide association results. Putative regulatory networks were constructed from 250 high-scoring pairs of proteases and related genes. This analysis suggested thirteen network modules, five diagnosis-based and eight biomarker-based. Through genetic associations and published literature on module members, the modules could be classified into different disease modalities including cholesterol homeostasis and high blood pressure.

## Introduction

Proteolytic enzymes, i.e. proteases, have been known to play key roles in biological processes such as cell cycle progression, tissue remodeling, neuronal outgrowth, hemostasis, wound healing, immunity, angiogenesis and apoptosis. ^1^ According to the latest data, more than 10% of the human protease genes are involved in human hereditary pathologies. ^2^ Several proteases have been validated as a drug targets, and are modulated by FDA-approved drugs like ACE inhibitors for hypertension, anti-PCSK9 monoclonal antibody for hyperlipidemia, DPP4 inhibitors for type II diabete mellitus, and so on. However, drug development efforts focused on proteases have been impaired due to lack of knowledge of the complex mechanisms by which their activities are regulated and of the multiple functions they have in diverse biological pathways. In addition, they are often linked into proteolytic cascades where cleavage by one protease activates multiple other proteases.

*MEROPS*^3^ is a manually curated database for a large number of proteases, their inhibitors, and substrates in multiple species. Recently, multiple protein-protein interaction (PPI) networks of proteases and their inhibitors were reconstructed based on MEROPS and PPI databases. ^4^ By validating the estimated networks through gene expression profiles from the GTEx database, ^5^ several potential new interactions between proteases and inhibitors were predicted. However, we are still far away from a comprehensive understanding of the contribution of the interaction of proteases and their inhibitors or substrates to human diseases.

The genome-wide association study (GWAS) is a powerful tool to understand the mechanisms of various human diseases in terms of genetics. The statistical power of GWAS analysis is strongly dependent on the number of samples used in the analysis, therefore, genetic and phenotype data aquiered from large cohorts are essential. UK Biobank^6^ (UKB), established in 2006 and one of the largest biobanks in the world, has been collecting data on a wide variety of human phenotypes such as medical history, blood test and urine test as well as comprehensive genetic data on the participants. Findings from GWAS of human diseases using UKB data has been used to elucidate new disease mechanisms.

In this study, we analyzed genetic and phenotypic data from UKB to predict the genetic similarities of putative pairs of proteases and their inhibitors or substrates. We performed a large number of GWAS using medical diagnoses and quantitative phenotypes focusing on SNPs related to proteases, their inhibitors and substrates, and identified phenome wide gene-level associations (PheWAS). This PheWAS analysis elucidated that genetic contributions of proteases are vastly different among diseases and quantitative phenotypes. Furthermore, we evaluated the phenotypic similarity between genes based on these identified associations and constructed networks from gene pairs showing strong similarity. This analysis predicted disease- or quantitative phenotype-specific networks of proteases, inhibitors, and substrates. This study could provide clues for understanding the mechanisms how proteases are linked with human phenotypes including diseases.

## Methods

### Gene selection

Data from the *MEROPS* database^3^ was downloaded and extracted into a local database. A map of UniProt^7^ ids to gene symbols (`HUMAN_9606_idmapping.dat.gz`) was downloaded from the UniProt website^8^ at Feb 11, 2020.

Lists of human proteases and their inhibitors in *MEROPS* was selected from the `domain` table. Human protease substrates in *MEROPS* were selected from the `substrate` table, excluding those exclusively annotated as due to signal sequence processing (`ECO:0000255`).

### SNP selection

We initially filtered out SNPs with an info score below 0.9 of info score in imputed genomic data in UK Biobank to obtain high quality SNPs. Functional SNPs were selected from the following four eQTL datasets: GTEx, ^5^ DICE, ^9^ eQTLGen, ^10^ and ImmuNexUT. ^11^ For GTEx, significant eQTLs in 53 human tissues were downloaded from GTEx website. For DICE, significant eQTLs in 15 human blood immune cell types were downloaded from DICE website. For eQTLGen, significant eQTLs of human blood was downloaded from eQTLGen website. For ImmuNexUT, eQTLs included all associations between SNPs and genes of 28 human blood immune cell types downloaded from the NBDC database (Research ID: hum0214.v4), and then significant SNPs with a p-value less than 1×10^−3^ were extracted as significant eQTLs. In addition, SNPs were annotated by vcftools using dbSNP, ^12^ and those predicted to produce missense, nonsense, frame-shift, or splice variants in target genes were extracted.

### Individuals

We performed sample QC and extracted British participants, as previously described in Dr. Neale’s laboratory^13^ with slight modification. Using QC data provided by UK Biobank, we filtered out samples that matched any of the following criteria: 1) heterozygosity is outlier or missing, 2) status of kinship inference is “exclusion”, 3) more than ten third-degree relatives, 4) samples not used for PCA calculation, 5) karyotypes other than XX or XY. We also excluded individuals that had withdrawn from the UK Biobank Project as of August 8, 2021. To define British samples, we calculated the sum of normalized deviation of the first 6 principal components (PCs) from individuals with British heritage using the following equation,

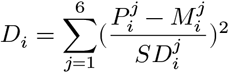

where *i* and *j* represent sample and PC number, respectively. *P* represents the provided PC score. *M* and *SD* the mean value and standard deviation of each PC calculated from samples that had “British” or “Irish” as their self-reported ethnicity, respectively. Samples with a *D*_i_ score less than 7^2^ were included in the final British population.

We did a PCA using the selected British samples to calculate new PC score for the association test. SNPs included in long-range LD regions^14^ were excluded. PCA was computed in Plink using SNPs with a MAF < 0.05 and r^2^ < 0.05.

### Phenotype data

#### Medical diagnoses

For medical diagnoses, we initially extracted ICD10 diagnosis codes in categories A00-R99 and Y40-Y49 for our samples. To account for disease with sex bias, the sex ratio of each ICD10 category was calculated, and ICD10 categories with a ratio exceeding 9:1 were classified into male and female-biased diseases, respectively. Other ICD10 categories were classified as unbiased diseases. ICD10 categories with less than 100 patients were removed.

#### Quantitative phenotypes

For 128 quantitative phenotypes, the original measured values in UK Biobank were used as provided. Estimated glomerular filtration rates (eGFR) were derived from serum creatinine and cystatin C levels using equations defined by CKD-EPI.^15^ The FIB-4 score to evaluate severity of liver disease was estimated from serum AST, ALT and platelet count.^16^ Ratio between the first second of forced expiration and the full forced vital capacity used in the diagnosis of lung disease was also calculated. Further, mean values of diastolic and systolic blood pressure and pulse rate from two measurements were calculated. To reduce measurement error, paired measurements with a difference exceeding the 90% quantile of all differences were excluded. In total, 135 quantitative phenotypes were obtained.

### Association test

Association tests for dichotomous and quantitative phenotypes were done by Firth logistic regression and linear regression in Plink2 v2.01,^17^ respectively. Sex, age, genotyping array, and the first five principal components were included as covariates. Female and male samples were removed in the association test of male- and female-biased ICD10 categories, respectively. P-values were converted to false discovery rate (FDR) by the Benjamini-Hochberg procedure.^18^ For each gene, the most significant SNP for each phenotype was used as a measure of gene-level association.

### Similarity index

To evaluate the phenotypic similarity among selected genes, rank biased overlap^19^ (RBO) was used as a similarity measure. For all combinations of proteases with all other selected genes, RBO was calculated based on the FDR in either medical diagnosis or quantitative phenotype GWAS results using the rbo function in the gespeR R package^20^ with a weight parameter of 0.7. Gene pairs having their transcription start sites (TSSs) within 10 Mbp of genomic distance in the same chromosome were removed. To extract gene pairs with high similarity, the threshold of RBO was set to 0.8 and 0.85 for medical diagnoses and quantitative phenotypes, respectively.

### Correlation network construction

Using gene pairs with high similarity, interaction networks were constructed. Interaction networks containing less than five genes were removed. For each network, the gene with the largest number of interactions was defined as the hub gene.

## Results

### Selection of genes, SNPs, individuals, and phenotypes

817 proteases, 324 protease inhibitors, and 4606 known protease substrates were extracted from the *MEROPS* database. As several genes belonged to multiple categories (Fig. 1), 5308 genes were selected in total. We further obtained SNPs involved in target genes from four human eQTL databases (GTEx,^5^ DICE,^9^ eQTLGen,^10^ and ImmuNexUT)^11^ and dbSNP (see method in detail). In total, 2,602,300 SNPs were selected for use in association tests. Using self-reported Biritish and Irish participant as a reference, 371,031 genetically British or Irish participants were selected in total.

**Figure 1:**
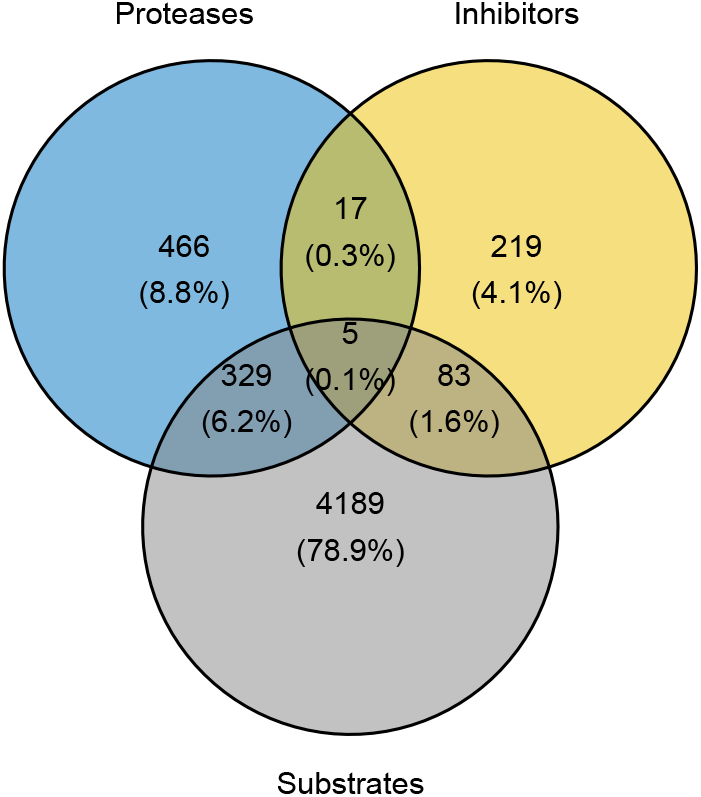
Venn diagram of gene categories.

Medical diagnoses were summarized on the ICD10 category level and analyzed as dichotomous phenotypes. To reduce sex bias, phenotypes were classified into three types, male-biased, female-biased and unbiased, based on their sex ratio (Table 1). We also obtained 131 quantitative phenotypes including 61 blood parameters, one arterial parameter, three parameters related to blood pressure, four urine parameters, two estimated glomerular filtration rates (eGFRs), one fibrosis score, four parameters evaluating lung function, 10 parameters related to visual function and 45 antigen tests. In total, we conducted genomic association tests for 1,405 phenotypes.

**Table 1:** Number of sex-biased medical diagnoses.

| Bias | ICD10 category |
| --- | --- |
| Unbiased | 722 |
| Female | 93 |
| Male | 21 |
| Total | 836 |

A complete list of medical and biomarker phenotypes can be found in Supplementary tables 1 and 2, respectively.

### PheWAS of medical diagnoses and quantitative phenotypes

Subsequently, we summarized the gene-level associations for each phenotype based on GWAS results to perform PheWAS analysis (see method in detail). As shown in Fig. 2 and Fig. 3, the number of genes that were significantly associated with each phenotype (BH-corrected FDR < 0.01) varied widely not only within diseases but also whithin quantitative phenotypes. In addition, we found that several phenotypes strongly associate with proteases. For example, the greatest numbers of disease-associated genes were found in the skin diseases Psoriasis (L40) and Vitiligo (L80) for all genes and proteases, respectively (Fig. 2). These results suggest that the genetic contribution of proteases and their target genes differs between human phenotypes.

**Figure 2:**
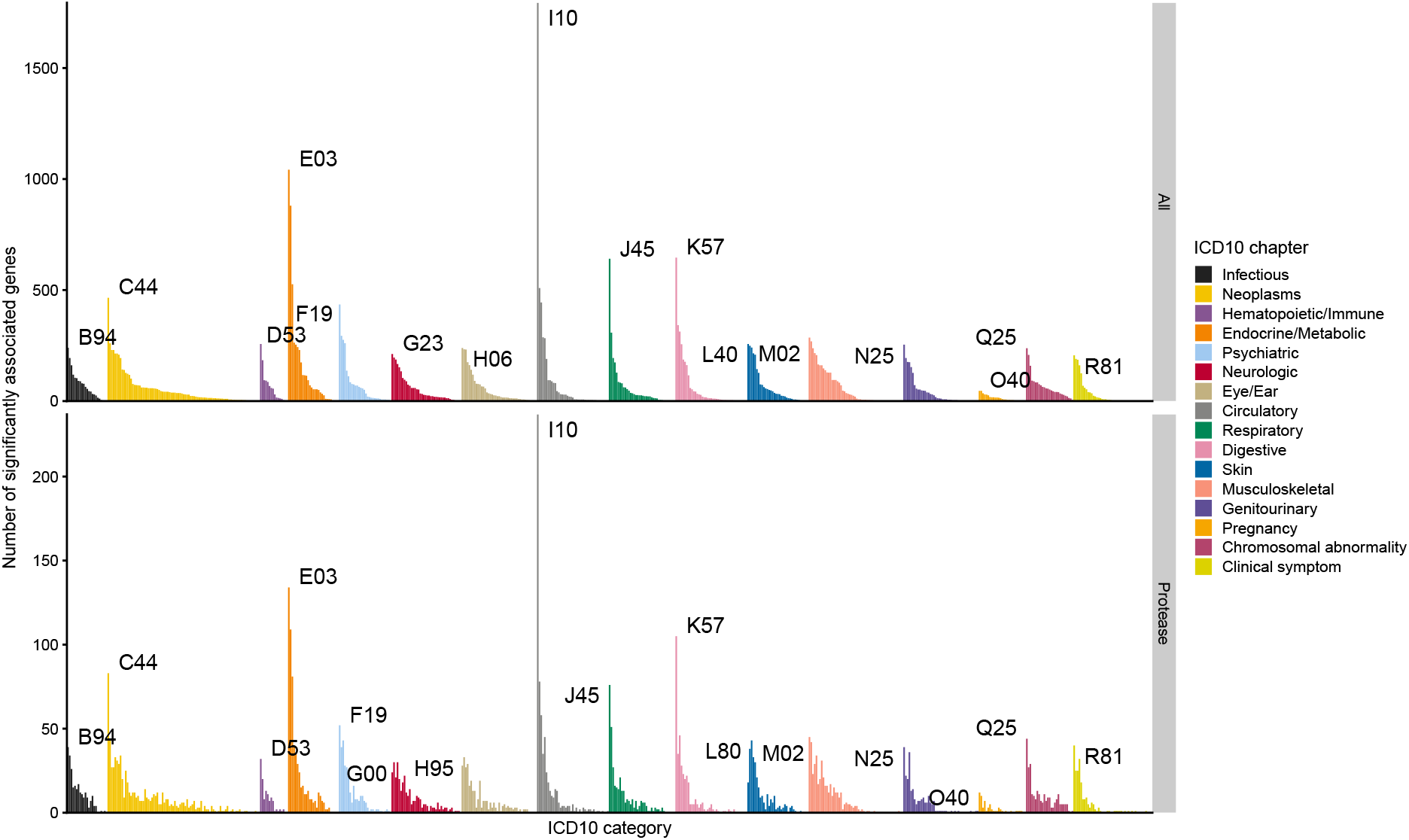
Number of genes showing significant associations to ICD10 categories. Genes with an FDR less than 0.01 were regarded as significant. Labels refer to the ICD10 category with the largest number of significant genes within each ICD10 chapter. The ICD10 categories were as follows: B94 Sequelae of other and unspecified infectious and parasitic diseases, C44 Other malignant neoplasms of skin, D53 Other nutritional anaemias, E03 Other hypothyroidism, F19 Mental and behavioural disorders due to multiple drug use and use of other psychoactive substances, G00 Bacterial meningitis, not elsewhere classified, G23 Other degenerative diseases of basal ganglia, H06 Disorders of lachrymal system and orbit in diseases classified elsewhere, H95 Postprocedural disorders of ear and mastoid process, not elsewhere classified, I10 Essential (primary) hypertension, J45 Asthma, K57 Diverticular disease of intestine, L40 Psoriasis, L80 Vitiligo, M02 Reactive arthropathies, N25 Disorders resulting from impaired renal tubular function, O40 Polyhydramnios, Q25 Congenital malformations of great arteries and R81 Glycosuria.

**Figure 3:**
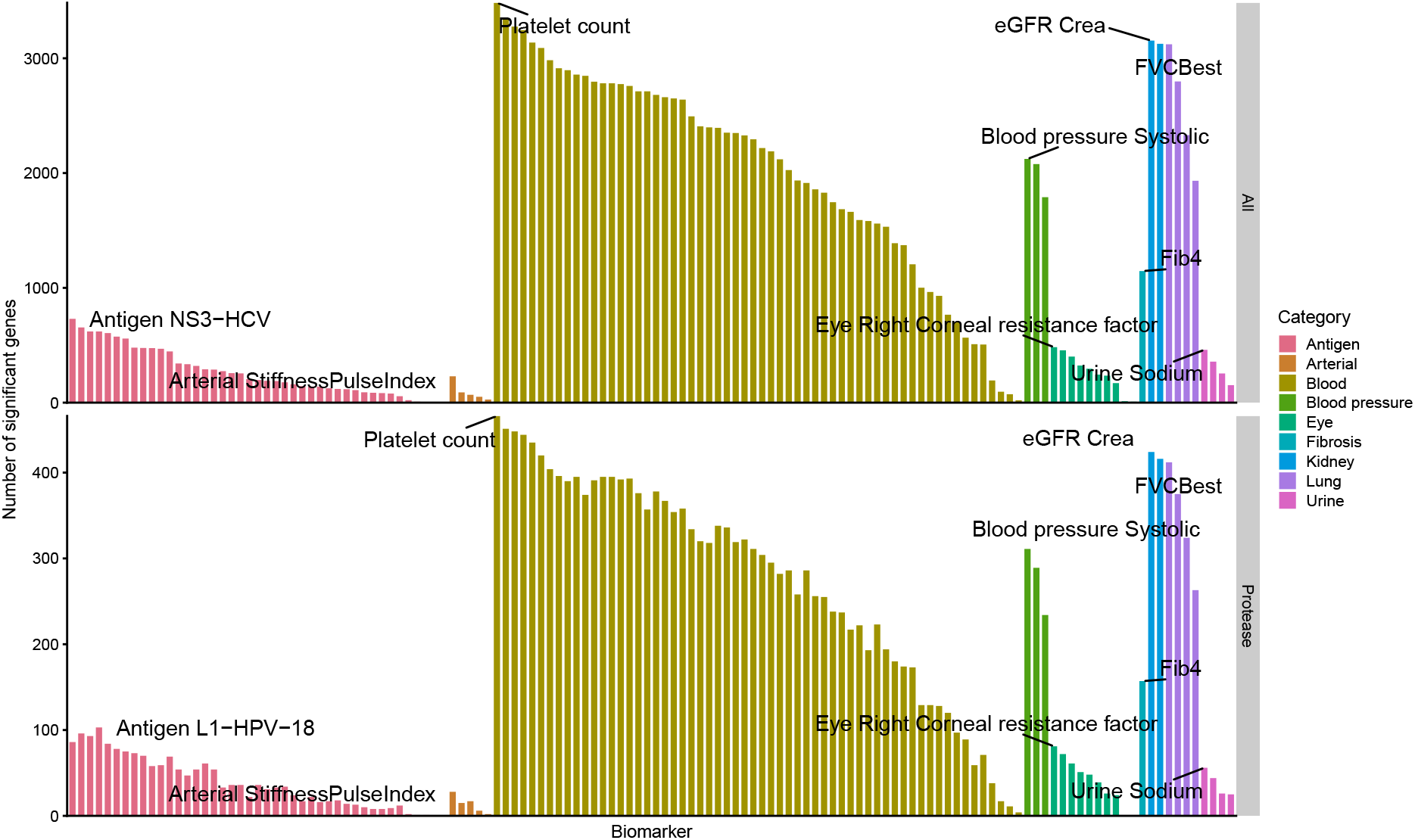
Number of genes showing significant associations to quantitative phenotypes. Genes with a FRD less than 0.01 were regarded as significant. Labels refer to quantitative phenotypes with the lagest number of significant genes per category.

### Phenotypic similarity

To explore the phenotypic similarity among proteases, their substrates and inhibitors, we used rank biased overlap (RBO) as a measure of similarity.^19^ For all combinations of 817 proteases and all 5308 genes, we calculated the RBO based on 836 medical diagnosis-based and 131 quantitative phenotype-based PheWAS results, and subsequently extracted gene pairs with an RBO greater than 0.8 and 0.85, respectively. To exclude similarities due to linkage disequilibrium, gene pairs with their transcription start site closer than 10 Mbp were excluded. This analysis identified 98 and 152 interacting gene pairs showing a similar association pattern of diseases and quantitative phenotypes, respectively (Fig. 4 and Table 2). The identified interactions contained a known interaction between TMPRSS6 and H2AC8 in quantitative phenotypes, whereas no known interaction was observed for medical diagnoses (Fig. 4). Moreover, we found no common interacting gene pairs between medical diagnoses and quantitative phenotypes (Table 2).

**Table 2:** Identified interactions between protease and target genes with top 5 RBO scores.

| PheWAS type | Protease | Target | RBO |
| --- | --- | --- | --- |
| Medical diagnosis | CELSR2 | TCP1 | 0.92 |
| Medical diagnosis | CELSR2 | IGF2R | 0.92 |
| Medical diagnosis | CELSR2 | ECSIT | 0.91 |
| Medical diagnosis | CELSR2 | ICAM1 | 0.91 |
| Medical diagnosis | PSMA5 | ICAM1 | 0.91 |
| Quantitative phenotype | TMPRSS6 | H2BC9 | 0.97 |
| Quantitative phenotype | TMPRSS6 | H2BC7 | 0.96 |
| Quantitative phenotype | TMPRSS6 | H4C5 | 0.95 |
| Quantitative phenotype | TMPRSS6 | H3C6 | 0.95 |
| Quantitative phenotype | TMPRSS6 | H2AC8 | 0.95 |

**Figure 4:**
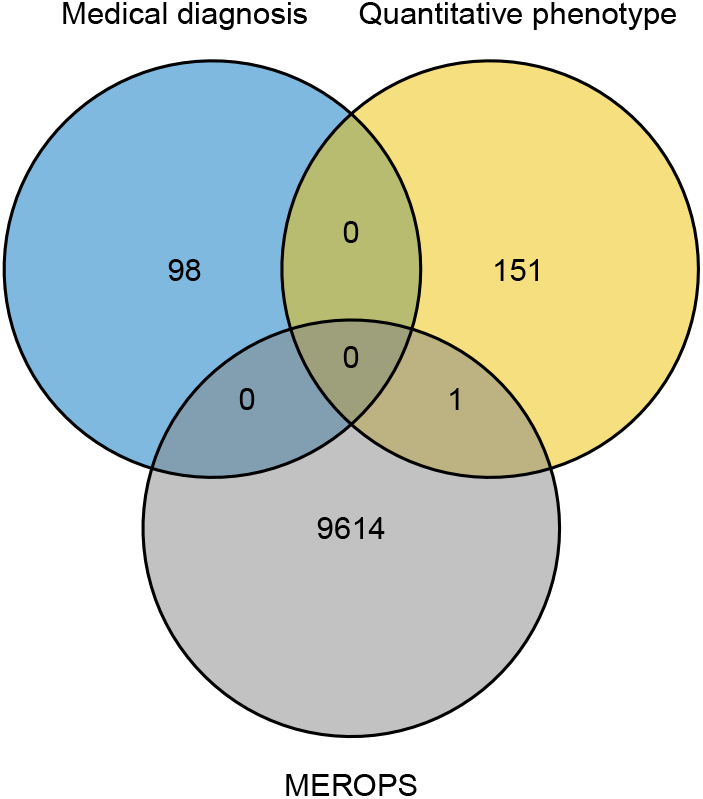
Venn diagram of interacting gene pairs identified using medical diagnoses and quantitative phenotype-based PheWAS.

A list of all suggested interactions can be found in Supplementary table 3.

To identify gene modules with similar phenotypic associations, we next constructed networks based on identified interactions. Using medical diagnosis-based PheWAS, we identified 5 modules, their hub gene, and their genetic association patterns (Fig. 5, Fig. 6, and Table 3).

**Table 3:** Summary of gene modules identified in medical diagnosis-based PheWAS.

| Module | Number of genes | Number of interactions | Number of known interactions |
| --- | --- | --- | --- |
| M1 | 16 | 36 | 0 |
| M2 | 9 | 12 | 0 |
| M3 | 9 | 9 | 0 |
| M4 | 8 | 9 | 0 |
| M5 | 5 | 4 | 0 |
| Total | 47 | 70 | 0 |

**Figure 5:**
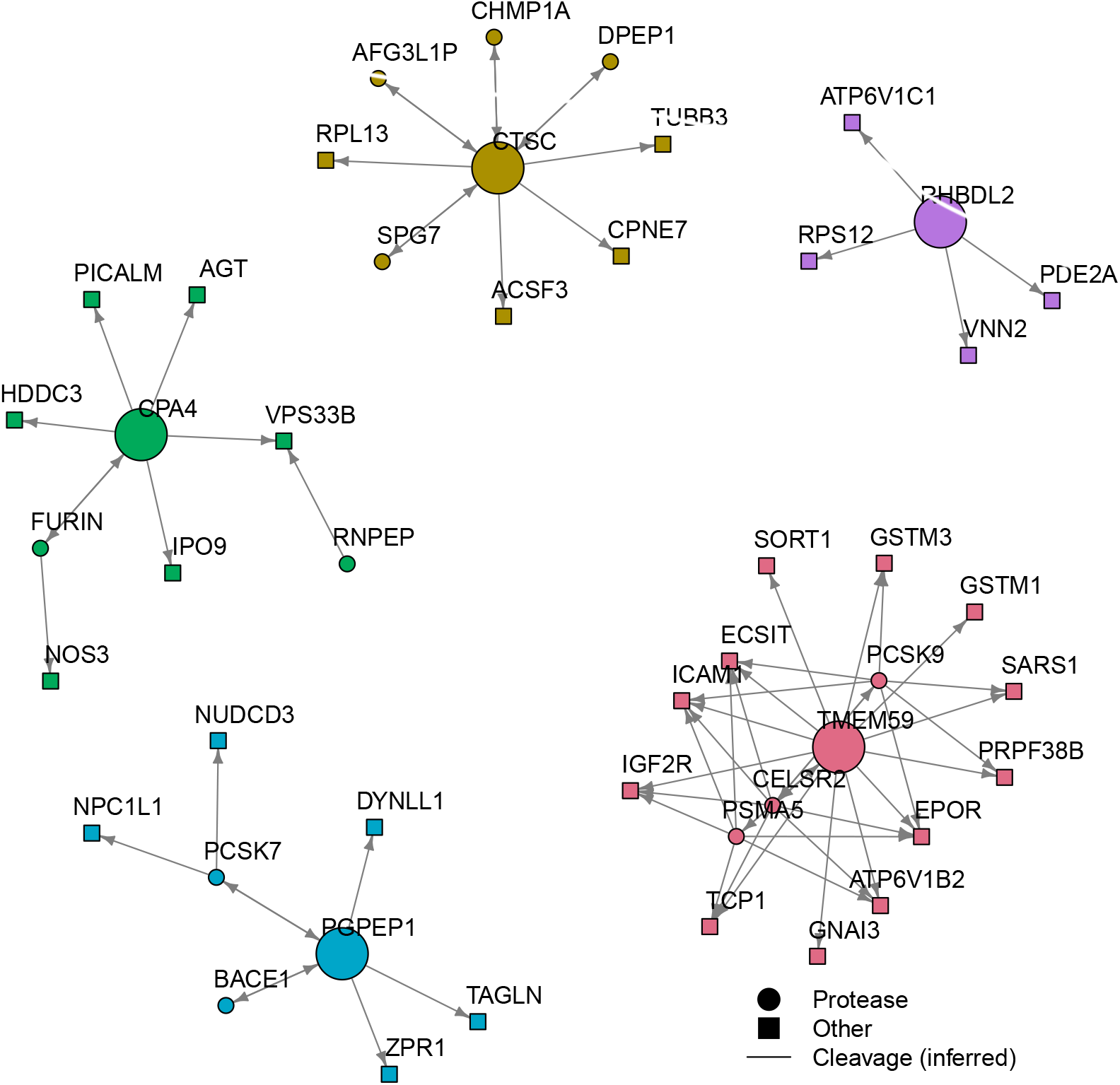
Gene module with similar association for diagnoses. The color and size of nodes represent the module and the hub gene, respectively.

**Figure 6:**
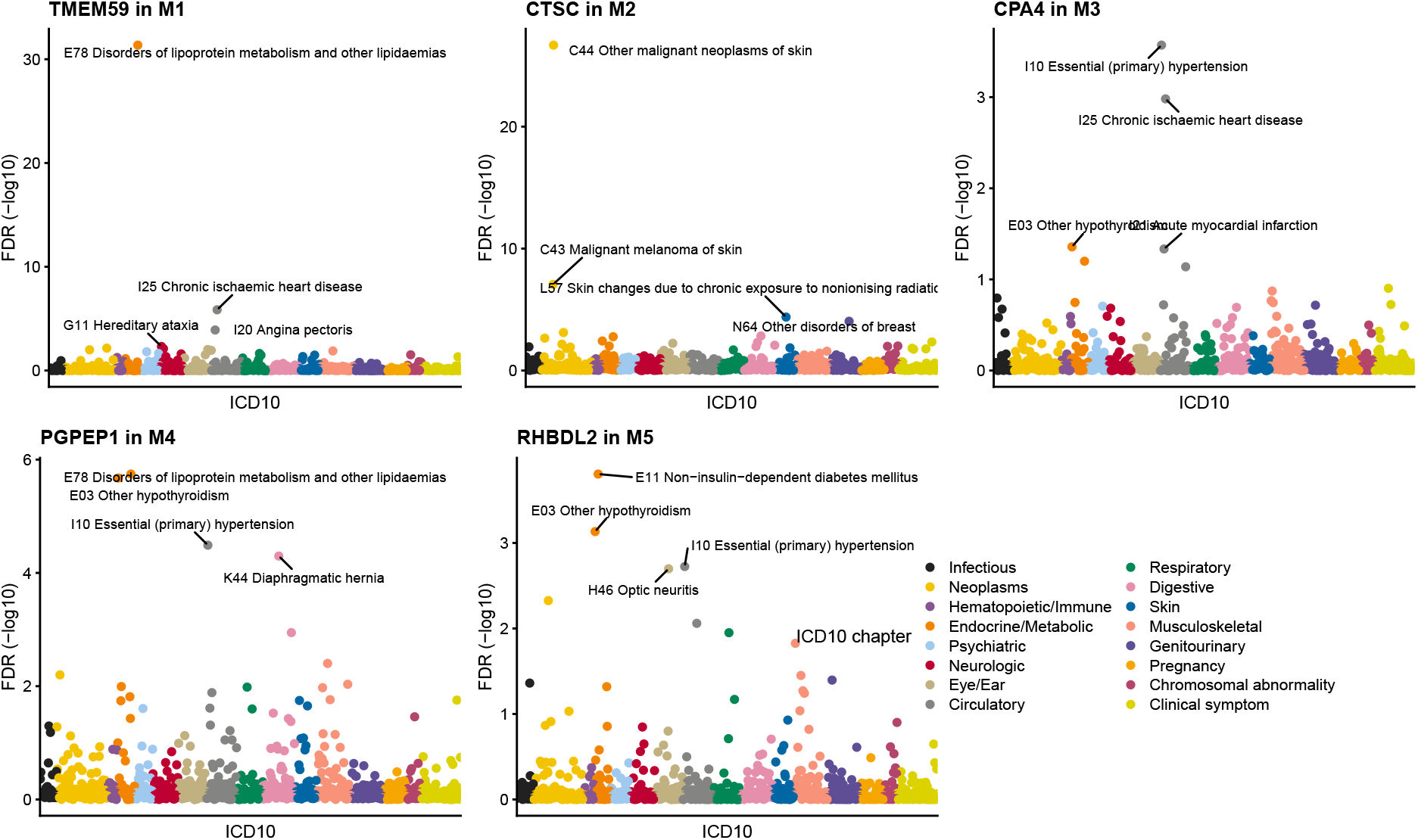
PheWAS of hub genes in each diagnosis module. Labels refer to ICD10 categories with top 4 FDR.

The largest gene cluster was M1 consisting of 16 genes. The hub gene, TMEM59 showed strong associations to Disorders of lipoprotein metabolism and other lipidemias (E78), Chronic ischemic heart disease (I25), Angina pectoris (I20), and Hereditary ataxia (G11), suggesting that genes in this module are involved with regulation of cholesterol homeostasis. Actually, we found several known regulators of cholesterol homeostasis such as PCSK9^21^ and SORT1.^22^ Similarly to M1, the M4 module consisting of eight genes was related to cholesterol homeostasis. The M4 hub gene PGPEP1 showed strong associations to Disorders of lipoprotein metabolism and other lipidemias (E78), Other hypothyroidism (E03), Essential (primary) hypertension (I10), and Diaphragmatic hernia (K44). This module also included several cholesterol regulators such as NPC1L1^23^ and TAGLN.^24^

The second largest gene clusters were M2 and M3, both of which included nine genes. The M2 hub gene CTSC (cathepsin C) had associations to Other malignant neoplasms of skin (C44), Malignant melanoma of skin (C43), Skin changes due to chronic exposure to non-ionizing radiation (L57), and Other disorders of breast (N64), suggesting involvement with skin diseases. On the other hand, the M3 hub gene CPA4 showed associations to Essential (primary) hypertension (I10), Chronic ischemic heart disease (I25), Other hypothyroidism (E03) and Acute myocardial infarction (I21), implying involvement with the regulation of blood pressure. AGT (angiotensin) and FURIN^25^ has been reported to be relevant with blood pressure regulation and hypertension, respectively.

Finally, the M5 module consisted of five genes, and the hub gene RHBDL2 showed strong associations to Non-insulin-dependent diabetes mellitus (E11), Other hypothyroidism (E03), Essential (primary) hypertension (I10) and Optic neuritis (H46). This module might be relevant to glucose metabolism and insulin signaling.

We further constructed correlation networks using interactions derived from quantitative phenotype-based PheWAS results, identifying eight modules, their hub genes, and their genetic association patterns (Fig. 7, Fig. 8, and Table 4).

**Table 4:**
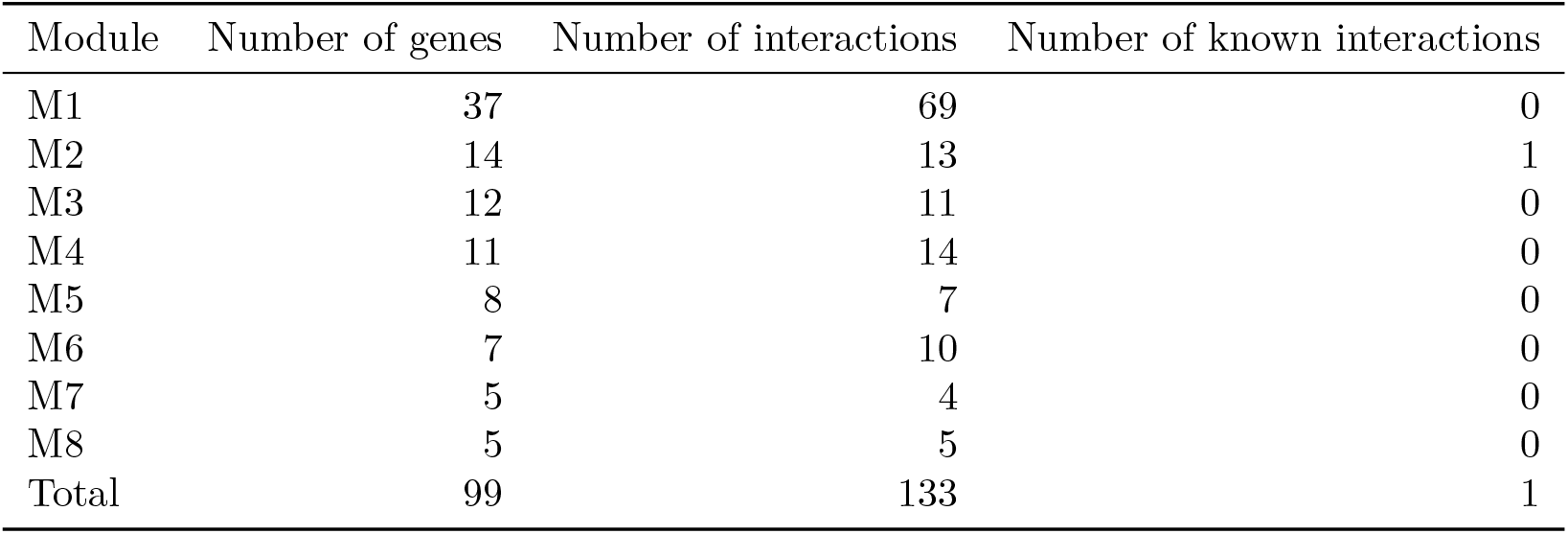
Summary of gene modules identified in quantitative phenotype-based PheWAS.

**Figure 7:**
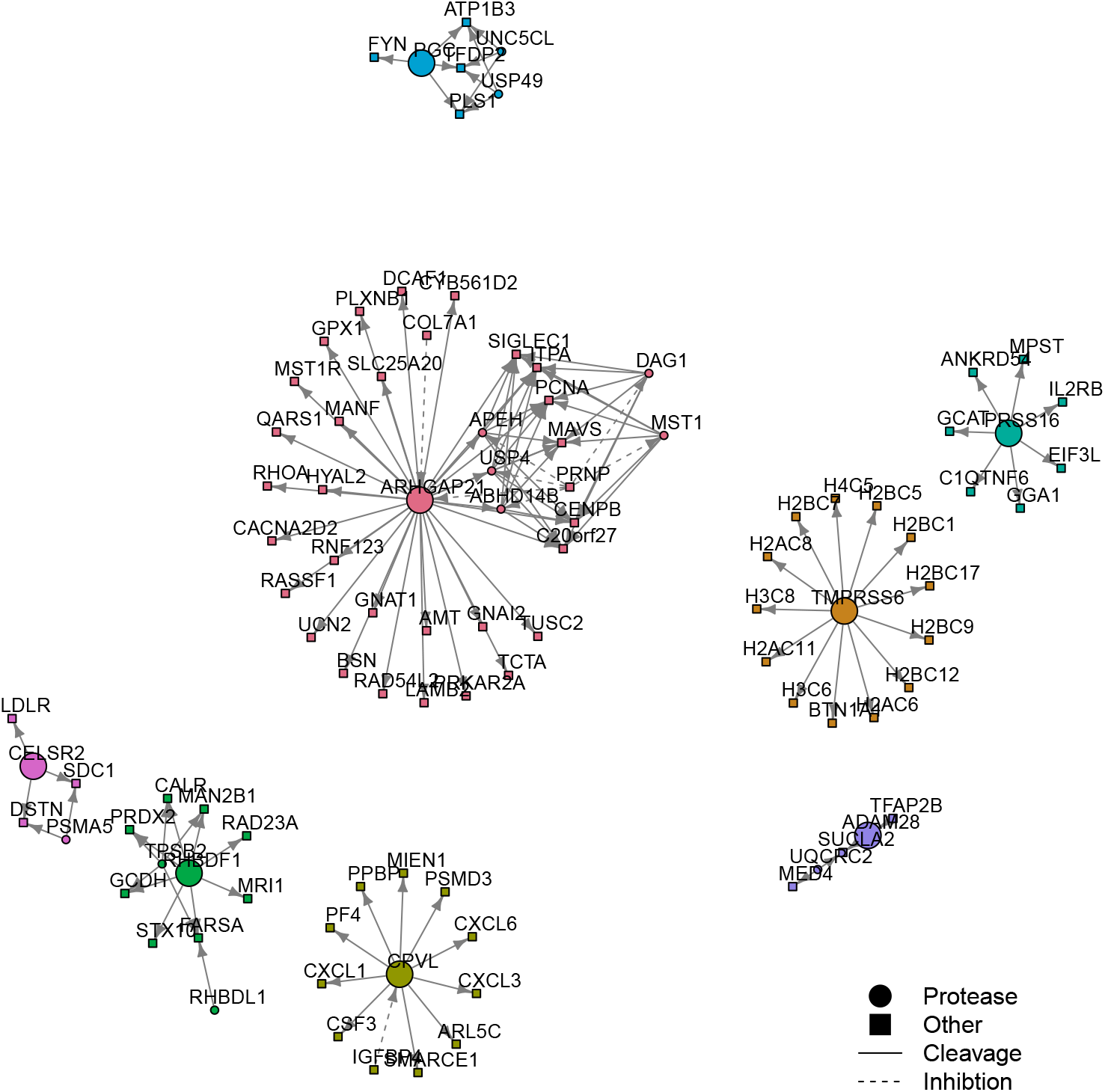
Gene module with similar association for quantitative phenotypes. The color and size of nodes represent the module and a hub gene, respectively.

**Figure 8:**
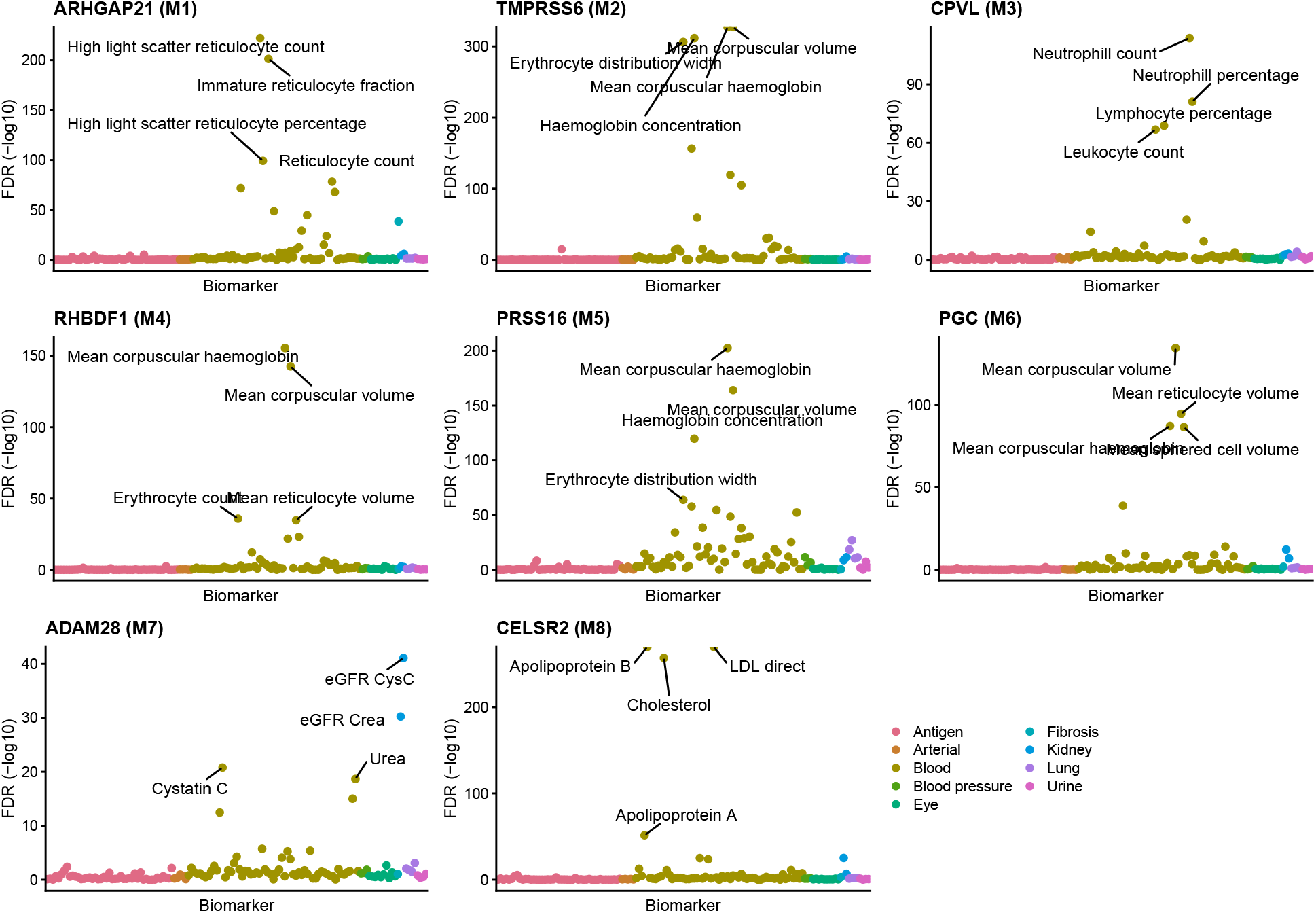
PheWAS of hub genes in each biomarker module. Labels refer to quantitative phenotype category with top 4 FDR.

The largest gene cluster was M1 consisting of 37 genes, in which the hub gene ARHGAP21 showed strong associations to reticulocyte-related phenotypes. This result implied that the M1 module contributes to the regulation of reticulocyte. In contrast to M1, the M2, M4, M5 and M6 modules showed associations for hemoglobin-related phenotypes. The third largest M3 module consisted of 12 genes, and the hub gene CPVL showed associations to leukocyte-related phenotypes. These five modules strongly associated with blood-related phenotypes.

By contrast, in the M7 module including five genes, the hub gene ADAM28 displayed associations to eGFR and serum urea levels. This suggests that the M7 module could be involved with renal function. In addition, a hub gene CELSR2 in the M8 module clearly showed associations to cholesterol-related phenotypes. This module included the LDL receptor (LDLR) regulating the uptake of LDL cholesterol.

Taken together, our PheWAS-based similarity analysis identified multiple functional protease modules which contribute to different human phenotypes such as disease, blood chemistry parameters and renal function. The interactions with proteases identified in this study were almost totally abstent from the *MEROPS* database, therefore further experimental validation is required.

## Discussion

To our knowledge, this study is the first that uses genetic associations for multiple human phenotypes to predict interactions between proteases and their inhibitors and substrates. Previous research has applied co-expression, co-regulation of promoters, to predict protease-inhibitor associations.^4^ However, since proteolytic cascades are very fast and often involve proteins expressed in distinct tissues and cell types, co-expression is not a suitable indicator of functional relatedness for proteases. Applying a conservative cut-off for our rank-based similarity measure we have predicted 98 and 152 interacting gene pairs from similar association pattern of medical diagnoses and quantitative phenotypes, respectively.

This set of interactions is almost completely absent from the *MEROPS* database of proteolytic events, making it difficult to assess the plausibility of these new interactions. We thus choose to use a literature search to evaluate the functional similarity of related genes. A graph-based analysis of the proposed interactions produced 13 gene modules, five derived from medical diagnoses and eight from quantitative phenotypes.

In the medical diagnosis-based analysis, M1 module contained the well-known cholesterol regulator PCSK9, which positively regulates serum low density lipoprotein (LDL) level through inhibiting the recycling of LDL receptors.^21^ In addition, SORT1 (sortilin 1) has been reported to be involved with cholesterol metabolism.^22^ In M4 module, NPC1L1 (Niemann-Pick C1-like 1) is a polytopic transmembrane transporter for cholesterol absorption in gastrointestinal tract epithelial and hepatic cells, and TAGLN (transgelin) is an actin-binding protein recently reported to regulate the endocytic uptake of LDL in HepG2 cell.^24^ In particular, the anti-PCSK9 monoclonal antibodies and a inhibitor of NPC1L1 have been approved by FDA and clinically used for treatment of hyperlipidemia. Cathepsin C, which is a hub gene in M2 module, has been reported to be upregulated in multiple squamous cancer cells including several sites of skin and to contribute to the development of squamous carcinogenesis.^26^ Furthermore, AGT (angiotensin), a well-known critical peptide hormone regulating blood pressure, was inferred to be cleaved by CPA4 in M3 module. Although this interaction was not included in *MEROPS* database, it was previously confirmed by *in vitro* experiments.^27^ In this module, FURIN has been also reported to be a potential serum biomarker for hypertension in the Chinese population.^25^

In the quantitative phenotype-based analysis, M1 module contains GPX1 (glutathione peroxidase-1), which is an important catalytic antioxidants in red blood cell. The serum level of GPX1 has been reported to negatively correlate with serum reticulocyte counts.^28^ The hub gene in M2, TMPRSS6, plays an important role in the hepcidin response to maintain iron homeostasis in the body,^29^ contributing to the regulation of red blood cells. Actually, multiple causal genetic mutations affecting TMPRSS6 enzymatic activity were found in iron-refractory iron deficiency anemia patients.^30^ In addition, FYN, a tyrosine kinase included in M6 module has been recently reported to modulate anti-oxidative response in red blood cell through the activation of glucose 6 phosphate dehydrogenase.^31^ CPVL (carboxypeptidase) in M3 module is localized at both the endoplasmic reticulum and membrane ruffles in primarily monocytic lineage cells including macrophages and potentially involved in the secretory pathway.^32^ This implied the influence of CPVL on the response of immune cells such as neutrophils and lymphocytes via cytokine secretion. Interestingly, multiple interactions between CPVL and chemokines such as CXCL1, CXCL3, CXCL6, and CSF3 were predicted in this module. In the M7 module associating with renal function phenotypes, an interaction between ADAM28 and TFAP2B was predicted. TFAP2B, known as AP-2, has been recently shown to regulate the differentiation of nephron progenitors in distal convoluted tubules, and its deficiency caused renal failure and fibrosis.^33^ M8 module included the LDL receptor regulating the uptake of LDL cholesterol into cells, supporting that the M8 module is linked to the regulation of cholesterol. CELSR2 and PSMA5 were found in both this module and hyperlipidemia-related M1 module in medical diagnosis-based analysis, even though no interactions were shared. Therefore, these genes might have strong genetic contribution specific to the regulation of cholesterol homeostasis, distinct from the other genes included in the same module.

## Conclusion

Proteases are involved in numerous biological processes but due to their complex regulation it is difficult to disentangle their functional involvement through molecular biology. In this project we tried to expand the understanding of how human proteases and their substrates are involved in human diseases by studying their connection to human phenotypes. By utilizing the large-scale genetic and phenotypic data from the participants in the UK Biobank we have produced data that is complementary to what can be achieved through cell- and animal-based studies. Putative regulatory networks were constructed from high-scoring pairs of proteases and related genes. This analysis suggested thirteen network modules, five diagnosis-based and eight biomarker-based. Through genetic associations and published literature on module members, the modules could be classified into different disease modalities including cholesterol homeostasis and high blood pressure.

## Author contributions

Conceptualization, T.E.; Data Curation, T.E.; Investigation, K.I. and T.E.; Methodology, K.I. and T.E.; Software, K.I. and T.E.; Visualization, K.I.; Writing, K.I. and T.E.; Resources, T.E.; Supervision., T.E.

## Acknowlegment

This research has been conducted using the UK Biobank^6^ Resource under Application Number 42755.

